# Identification and functional characterization of novel murine DIS3L2 isoforms

**DOI:** 10.64898/2026.10.22.761083

**Authors:** Leidy Paola P. Cepeda, Bruno R. S. Queiroz, Katherine Tsantarlis, Valdir Gomes Neto, Luiz H. S. Maniero, Bernadina A. Uscata, Hiro Goto, Carla C. Oliveira

**Author notes:** These authors contributed equally to this work (Leidy Paola P. Cepeda, Bruno R. S. Queiroz).

## Abstract

RNases are key factors for the posttranscriptional control of gene expression due to their involvement in processing, degradation and quality control of all classes of RNA in eukaryotes. DIS3L2 is an evolutionarily conserved exoribonuclease involved in cytoplasmic RNA degradation pathways that regulate gene expression. Macrophages undergo great change in gene expression of specific genes that allow these cells to perform multiple functions in addition to defense against infection. Here, we analyzed the change in expression of macrophages genes in response to *Leishmania major* infection, and show evidence that the levels of RNases such as the RNA exosome, XRN1, and DIS3L2, are modulated by the infection. We then identified three DIS3L2 variants that are differentially expressed in murine macrophages and other cell lines. DIS3L2 isoforms show similar RNA binding affinity, but differences in exonucleolytic activity in *in vitro* assays. Furthermore, we characterized the interactome of the DIS3L2 variants, which shows similarities with respect to ribosomal proteins, but differences that hint at specific functions for each isoform. This work uncovers previously uncharacterized DIS3L2 isoforms that are active, whose functions have yet to be identified.

## Introduction

RNases are very important for the posttranscriptional control of gene expression due to their involvement in processing, degradation and quality control of all classes of RNA in eukaryotes. Exo- and endoribonucleases, also control expression of individual mRNAs and small regulatory RNAs, thereby affecting transcription and translation [1–4].

Among the eukaryotic exonucleases, the RNA exosome participates in the degradation of cytoplasmic mRNAs, directly regulating transcripts levels [5–8]. In mammals, the RNA exosome complex is composed of ten different protein subunits, nine of which form the catalytically inactive *core* (Exo9 – subunits EXOSC1-9), present in the nucleus and cytoplasm, that associates with the active exonucleolytic RNases, the nucleolar EXOSC10, the nucleoplasmic DIS3, or the cytoplasmic DIS3L [9]. The paralog enzymes DIS3 and DIS3L belong to the RNase II/RNB family of exoribonucleases, have functional RNB domains and display processive exonuclease activity. In addition, both enzymes contain the endonuclease PIN domain, but this is inactive in DIS3L [10–12].

In addition to these two DIS3 paralogs, mammalian cells have a third cytoplasm-exclusive paralog, DIS3L2, an evolutionarily conserved protein that retains the RNB domain, but lacks the PIN domain and has been shown to be involved in cytoplasmic mRNA degradation, independently of the exosome, and to preferentially degrade uridylated RNAs [13–16]. DIS3L2 also targets ARE-containing transcripts and has been shown to be involved in the regulation of various cellular processes, including rRNA biogenesis, endoplasmic reticulum-associated mRNA translation, cell differentiation and proliferation, T-cell activation, spermatogenesis, and nonsense-mediated mRNA decay (NMD) [10, 13, 17]. The structure of mouse DIS3L2 with a poly-U substrate has been determined, which shows that its three RNA binding domains form a funnel leading the substrate towards the catalytic site through a path different from the yeast ortholog exosome-associated Rrp44/Dis3 [10, 17, 18].

Gene expression of macrophages has been shown to be altered when these immune response cells are infected with intracellular pathogens such as *Leishmania sp.* [19, 20]. RNA-seq analysis of host-parasite interaction show that upon *L. major* infection, macrophage most substantial response at the gene expression level occurs early after infection, decreasing over time [21, 22]. These changes in gene expression are considered to occur at the transcriptional level, based on the analysis of RNA steady-state levels by microarray or RNA-seq. However, it remains to be investigated whether the lower mRNA levels could also be caused by RNA degradation.

In this work, we set out to investigate the role of murine macrophage RNases in posttranscriptional control of gene expression. We describe the identification of DIS3L2 isoforms expressed in these cells, and the analysis of the RNase activity of the purified recombinant DIS3L2 isoforms, demonstrating that all of them bind RNA and have exonuclease activity *in vitro*. The results shown here highlight the importance of DIS3L2 for the cytoplasmic control of gene expression in macrophages.

## Material and Methods

### Leishmania culture

*Leishmania major* (MRHO/Su/59/P) was maintained by regular passages in BALB/c mice, according to protocols approved by the Ethics Committee on Animal Use of the Faculdade de Medicina, Universidade de São Paulo (CEUA-FMUSP No. 086/14 and 087/14), and promastigotes were derived from amastigotes recovered from these animals. Promastigotes were cultured at 26 °C in M199 medium (Gibco™) supplemented with 10% heat-inactivated FBS (Cripion), 100 U/mL penicillin and 100 µg/mL streptomycin. Cultures were initiated at 1 × 10⁶ promastigotes/mL and grown until the stationary phase, when the parasites were used for infection.

### Cell culture

RAW 264.7 (macrophages; ATCC TIB-71), P3X63Ag8U.1 (P3U1, myeloma; ATCC CRL-1597) and L929 (fibroblasts; ATCC CCL-1) murine cell lines were cultured in 75 cm² culture flasks in RPMI-1640 (RAW 264.7) or DMEM (Dulbecco’s Modified Eagle’s Medium) (Gibco™) (P3U1 and L929) supplemented with 10% heat-inactivated fetal bovine serum, 100 U/mL penicillin and 100 µg/mL streptomycin, at 37 °C with 5% CO₂. The medium was replaced every two days, and cells were subcultured at ∼80% confluence, by gentle scraping (RAW 264.7) or using Trypsin-EDTA 0.05% (Gibco™) (P3U1 and L929).

### Macrophage infection

RAW 264.7 cells were seeded at 5 × 10⁵ cells/well in 24-well plates in the same medium and allowed to adhere for 1 h before infection. Cells were then infected with stationary-phase *L. major* promastigotes (8:1 parasite-to-macrophage ratio). After 6 h, non-internalized parasites were removed by washing with PBS, and cultures were maintained for 6 to 48 h post-infection. Uninfected cells cultured in parallel served as controls.

### RNA Isolation and cDNA synthesis

Total RNA was isolated from RAW264.7, P3U1 and L929 cells using TRIzol^®^ reagent (Thermo Fisher Scientific) according to the manufacturer’s protocol. Cells in 80% confluence were washed twice with ice-cold PBS and lysed directly using 1 mL of TRIzol reagent. Samples were incubated for 5 min at room temperature followed by the addition of chloroform (200 µL per mL of TRIzol). Centrifugation was performed at 12,000 × g for 15 min at 4 °C. The upper aqueous phase was mixed with isopropanol and subsequently centrifuged, washed with 75% ethanol, and then suspended in RNase-free water. RNA integrity was assessed by 1% agarose gel electrophoresis (visualization of 28S and 18S ribosomal bands). The RNA samples were treated with RNase-free DNase I (Invitrogen) at 37 °C for 30 min. First-strand cDNA was synthesized from 2 µg of total RNA using the SuperScript III First-Strand Synthesis System (Invitrogen) according to the manufacturer’s instructions.

### PCR amplification of different DIS3L2 isoforms

PCR amplification was performed using Phusion High-Fidelity DNA Polymerase (Thermo Scientific) according to the manufacturer’s instructions. Each 50 µL reaction contained 1× Phusion HF Buffer (1.5 mM MgCl₂), 200 µM dNTP, 0.5 µM forward and reverse primers (Suppl. Table 1), 1 µL of 2X-diluted cDNA template, and 0.02 U/µL Phusion DNA Polymerase; the final volume was adjusted to 50 µL with nuclease-free water. PCR products were resolved by electrophoresis on a 2% agarose gel containing ethidium bromide and visualized under UV illumination.

### Quantitative real-time PCR

Gene expression levels were quantified by real-time quantitative PCR (RT-qPCR) using HOT FIREPol® EvaGreen® qPCR Supermix (5×) (Solis BioDyne). Each reaction (final volume of 10 µL contained 1× HOT FIREPol® EvaGreen® qPCR Supermix, 0.2 µM of each forward and reverse primer (Suppl. Table 2), and 2 µL of diluted cDNA template. Reactions were performed in 96-well plates on a 7500 Real-Time PCR System (Applied Biosystems). Cycling conditions consisted of an initial hot-start activation step at 95 °C for 12 min, followed by 40 cycles of denaturation at 95 °C for 15 s, annealing at 60 °C for 30 s and extension at 72 °C for 30 s. A melting-curve analysis (60–95 °C, 0.5 °C increments) was performed as the instrument’s default settings at the end of each run to confirm amplicon specificity. CYC1, Actin, and GAPDH genes were used as internal references [23, 24]. −ΔCt values were standardized independently within each cell line. For each cell line, the mean (μ) and standard deviation (σ) were calculated across all analyzed DIS3L2 splice variants in each cell line, and Z-scores were obtained as Z=[(−ΔCt)−μ]/σ [25]. Thus, Z-score values represent the relative position of each splice variant with respect to the overall distribution of DIS3L2 splice-variant abundance within the corresponding cell line. Heatmap was used for visualization of the data.

### Recombinant DIS3L2 isoforms purification

The DIS3L2 isoform genes were synthesized and cloned into pET-28a by BioLinker Biologia Sintetica; the constructs have a 6xHis tag and MBP at the N-terminus of the DIS3L2 protein. His-MBP DIS3L2 was expressed in Rosetta pLYS (DE3) cells. Cells were grown at 37 °C to an optical density of 0.4 to 0.6 in LB, and protein expression was induced by adding 0.1 mM IPTG at 18 °C overnight. The cells were harvested and pellets stored at −80 °C until purification. Cells were lysed in Avestin Emulsiflex C3 in 50 mM HEPES pH 8.0, 500 mM NaCl, 10% glycerol, 5 mM MgCl_2_, 10 mM imidazole, and 5 mM BME, with a complete EDTA-free protease inhibitor tablet (Roche). Purification was conducted using HisTrap FF Crude 5 mL (Cytiva). The resin was washed with lysis buffer and then with 50 mM HEPES pH 8.0, 250 mM NaCl, 10% glycerol, 5 mM MgCl2, 50 mM imidazole, and 5 mM BME. Samples were eluted in the same buffer, with the addition of 250 mM Imidazole. The protein eluate was cleaved with TEV protease to release the 6xHis-MBP tag and loaded onto a SEC column (HiLoad 26/600 Superdex 200 PG), equilibrated with 50 mM HEPES pH 8.0, 250 mM NaCl, 5% glycerol, 5 mM MgCl_2_, and 5 mM BME. The final protein concentration was determined using a spectrophotometer. To ensure the protein preparation was RNA-free, A_260_/A_280_ ratio was measured and values varied between 0.5 and 0.6. The DIS3L2 protein concentration was calculated using the Beer-Lambert equation with an extinction coefficient estimated using Expasy ProtParam protein characterization tools (https://web.expasy.org/protparam/).

### RT-FeDEx Assay

The purified DIS3L2 isoforms (100 nM) were incubated at 30 °C in the presence of 0.3125 pmol μL^−1^ 5′-FAM-RNA and 0.625 pmol μL^−1^ DNA-TAMRA-3′ in reaction buffer containing 50 mM Tris-HCl (pH 8.0), 2 mM MgCl_2_, 50 mM NH_4_Cl, and 0.5 mM DTT [26, 27]. FAM-RNA and DNA-TAMRA were heated at 65 °C prior to assays in order to allow the formation of hybrid molecules. RNase A was used as a positive control for enzyme activity. Samples containing only RNA or the hybrid molecule RNA/DNA with no protein sample added were used as negative controls. Fluorescence emission was detected on the 7500 Real-Time PCR System (Applied Biosystems). The final degradation products were resolved on a 20% polyacrylamide gel in 1X TBE, and images were acquired on a ChemiDoc MP equipment (Bio-Rad). For FAM detection, a fluorescein filter 460 nm of emission and excitation 518 nm. TAMRA signal was detected with the Rhodamine filter, with emission at 520 nm and excitation at 577 nm.

### EMSA

Gel electrophoresis mobility shift assay (EMSA) was used to detect DIS3L2 complex with RNA. Different concentrations (from 6.2 to 800 mM) of purified recombinant DIS3L2 variants were incubated with 1 pmol of single strand 5’-FAM-RNA for 30 min at 30°C in 50 mM Tris-HCl (pH 8.0), 30 mM NaCl, 10 mM K_2_HPO_4_, 0.1 mM DTT, 0.1 mg/mL BSA, 10 mM heparin, 10 mM EDTA buffer and subsequently subjected to electrophoresis on a 12% native polyacrylamide gel in 1x TBE buffer. Bands were visualized on a ChemiDoc MP equipment (Bio-Rad).

### DIS3L2 Isoforms Pull-Down

The purified 6xHis-MBP-DIS3L2 isoforms were immobilized in Amylose resin (NEB), and total protein extract from macrophage RAW 264.7 was incubated with the resin with and without bound DIS3L2 isoforms in biological triplicates. The resin was washed, and the bound protein was eluted with 10 mM maltose according to the manufacturer’s protocol.

### LC-MS/MS analysis

The eluted proteins from the pull-down were quantified, denatured with 4 M urea and 5 mM DTT, alkylated with 14 mM iodoacetamide, and digested with trypsin for 16 h. The digestion was quenched with 0.4% TFA, and the tryptic peptides were desalted using the Pierce C18 tips according to the manufacturer’s protocol. Peptides were analyzed using a Thermo Fisher Scientific Orbitrap Exploris 240 mass spectrometer (Thermo Fisher Scientific) interfaced with an Evosep One liquid chromatography system via a nanoelectrospray ion source in the DIA mode. The peptides were preconcentrated on EV2001 C18 Evotips and separated on an EV1106 column (15 cm × 150 μm, 1.9 μm particle size) using the Evosep 30SPD method (30 samples per day). The nanoelectrospray voltage was maintained at 1.7 kV, and the source temperature was set to 275°C. The Orbitrap Exploris 240 was operated at a full MS resolution of 120,000, scanning across a mass-to-charge (m/z) range of 350–1400, with an automatic gain control (AGC) target of 300% and a maximum injection time 45ms. Tandem mass spectrometry (MS/MS) scans were conducted with 46 scan windows, each spanning 14.7 m/z units, from 361 to 1033 m/z, with an AGC target of 1000%, a normalized collision energy (NCE) of 27%, and a resolution of 15,000 [28–30]. Biological Process and Molecular Function were retrieved directly from the Gene Ontology resource and subsequently consolidated into higher-order categories for interpretation. Identifier for the raw mass spectrometry data: PXD083862, available via PRIDE.

### Polysome-Seq

The infected 6h post-infection (6 hpi) and non-infected macrophages were treated with cycloheximide (100 μg/mL) 5 min before harvest. The treated cells were lysed in hypotonic lysis buffer containing 5 mM Tris-HCl (pH 7.5), 2.5 mM MgCl2, 1.5 mM KCl, 100 mg/ml cycloheximide, 2 mM DTT, 0.5% Triton X-100, and 0.5% sodium deoxycholate, and cell debris was removed by centrifugation at 13,000 rpm for 2 min at 4 C. Lysates were loaded onto 7% to 47% sucrose density gradients (20 mM HEPES-KOH [pH 7.6], 100 mM KCl, 5 mM MgCl_2_) and centrifuged in a HITACHI CP80-NX, RP40-ST rotor at 39,000 rpm for 2.5h at 4°C. After centrifugation, the gradients were fractionated (500 μL per fraction), and the absorbance at 254 nm was continuously monitored. The RNA was extracted from polysome fractions using the phenol:chloroform method. The mRNA library was prepared with the SureSelect Strand-Specific RNA Library Prep System kit, and the library was sequenced on the Illumina HiSeq 2500 system. The adaptor and index sequences were removed from the raw reads using the Trimmomatic algorithm, aligned with the reference genome in HISAT2, and the differential expression was calculated in DESeq2. All tools were used via the Galaxy Europe server. The RNA-seq data generated in this study have been submitted to the NCBI Sequence Read Archive (SRA) under BioProject accession number PRJNA1537628.

## Results

Based on the description of gene expression changes in macrophages upon infection with intracellular pathogens, to test the hypothesis that these changes may occur at the posttranscriptional level, we infected RAW 264.7 cells with *Leishmania major* promastigotes at the ratio of 8:1 (*L. major*/macrophage) for 6 hours for analysis by RNA-seq experiments with samples of total RNA, or RNA isolated from translating polysomes, obtained from three biological replicates, and comparing with uninfected cells. The analysis of total RNA shows that the host responds to the infection by changing gene expression, repressing or activating different groups of genes (Fig. 1A; Suppl. Table 1). Interestingly, when analyzing the genes of RNases, such as the exosome subunits, we see some changes at the transcripts levels, most of them showing lower levels upon infection, with the exceptions of DIS3 and EXOSC10, which are upregulated at 6 hours post infection (hpi) (Fig. 1A). These subunits are catalytically active and concentrated in the nucleus, and mainly involved in pre-rRNA processing and RNA quality control [9, 31, 32]. Surprisingly, the analysis of mRNAs engaged in translation showed that while most of the exosome subunits showed lower mRNA levels at 6 hpi, the mRNAs of EXOSC10, DIS3 and its paralogs DIS3L and DIS3L2 had higher concentrations on polysomes (Fig. 1B).

**Figure 1.**
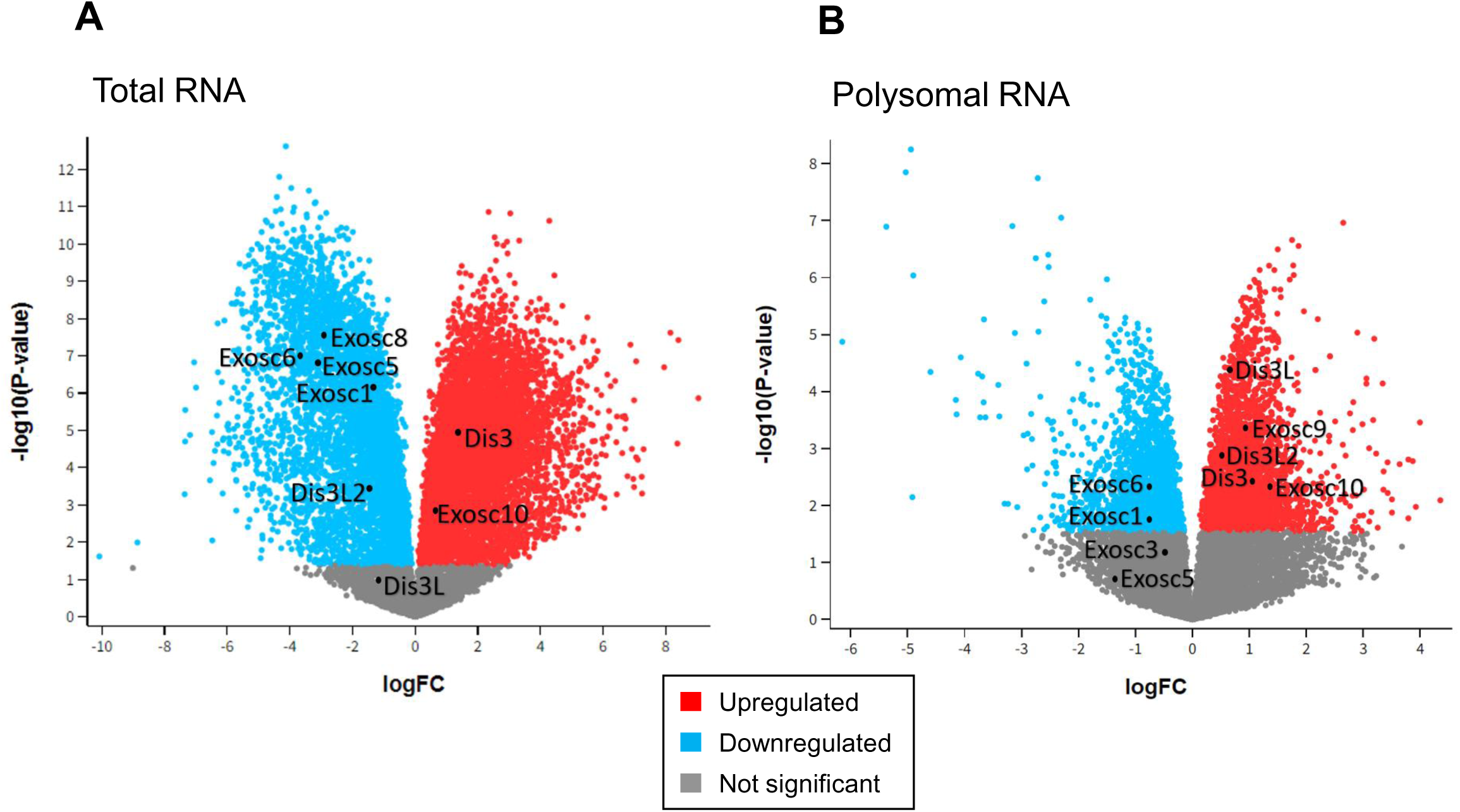
Transcriptomics analysis by RNA-Seq of *Leishmania major* infected macrophages at 6 hpi. Volcano plot comparing the two conditions (Infected vs Not Infected) indicating the different groups of genes altered in response to the infection with *L. major.* (**A**) Total RNA (**B**) RNA isolated from polysomal fractions of sucrose gradients.

More detailed analysis of the mRNAs of the exosome subunits that are present in polysomal fractions, and therefore engaged in translation, shows that the levels of expression of the catalytically active subunits, EXOSC10, DIS3 and DIS3L, increase at 6 hpi (Fig. 2A). On the other hand, most of the exosome core subunits EXOSC1-9, which are important for the recruitment and activity control of EXOSC10, DIS3 and DIS3L, show lower levels of expression at 6 hpi (Fig. 2A). The cytoplasmic exclusive DIS3 paralog DIS3L2, which is independent of the exosome, also showed higher levels in polysomes, indicating higher translational rate (Fig. 2A). Other cytoplasmic proteins involved in RNA degradation, such as the SKI complex subunits SKIC2, SKIC3 and SKIC7/HBS1L were also analyzed and had their levels increased at 6 hpi (Fig. 2B). Although SKIC8 shows lower levels of expression in total RNA, slightly higher levels of this mRNA are engaged in translation (Fig. 2B). The 5’-3’ exonucleases XRN1 and XRN2 also show higher levels at 6 hpi (Fig. 2B). Interestingly, transcripts of endonuclease subunits responsible for tRNA splicing (Tsen2,15,34,54) showed lower levels both in total RNA as in polysomes, indicating lower expression, and therefore, inhibition of translation (Fig. 2C). In agreement with this observation, translation initiation factor eIF2α kinases Eif2ak1-4 showed higher levels of total mRNA and in polysomes (with the exception of Eif2ak1), indicating translation repression at 6 hpi (Fig. 2C). These results suggest that *L. major* infection leads to a change in the levels of enzymes involved in regulation of posttranscriptional control of gene expression, consistent with an early cell response to the presence of the parasite.

**Figure 2.**
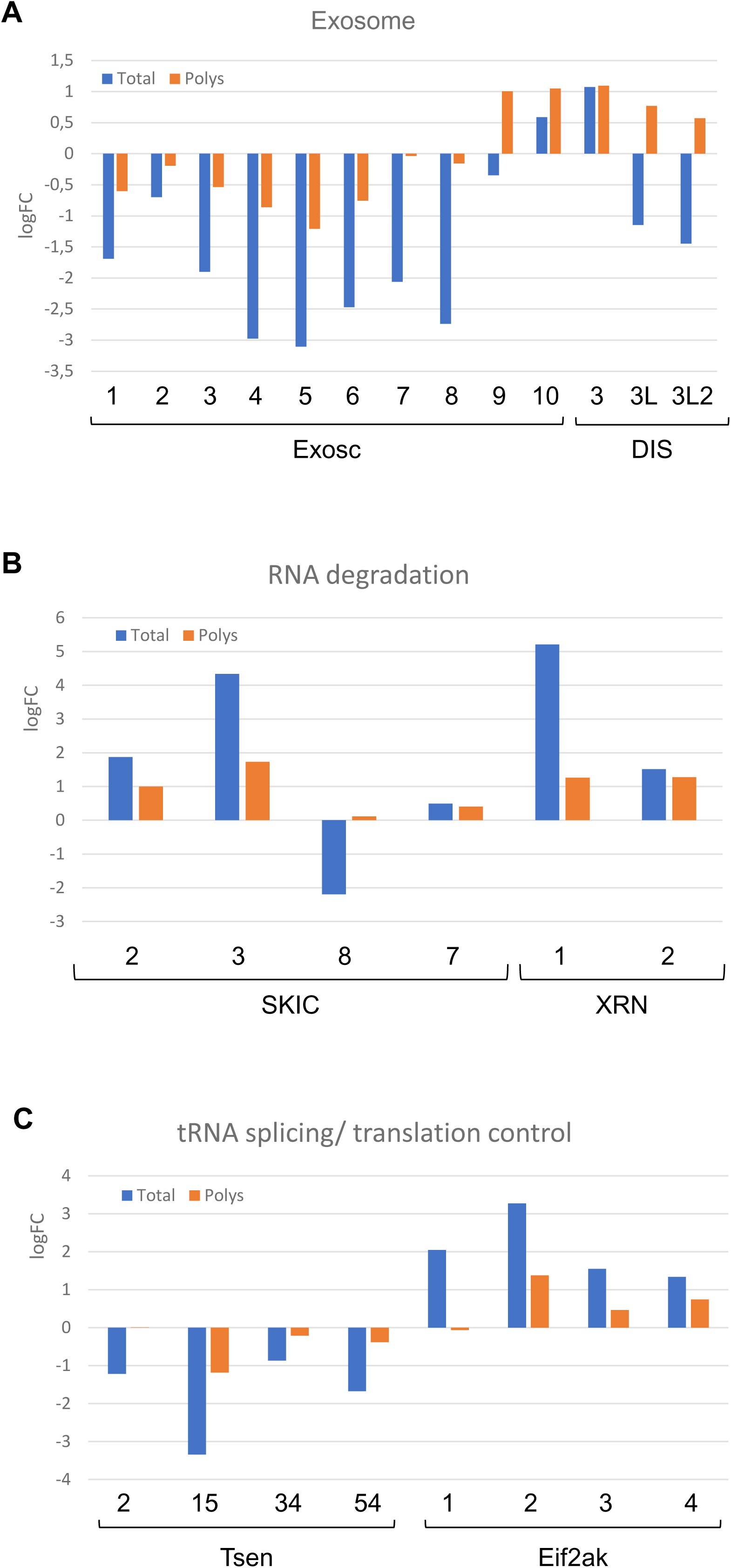
*L. major* infection leads to changes in expression levels of genes involved in mRNA processing. Graph showing RNA seq results with the altered expression levels of genes at 6 hpi, compared to uninfected cells. (**A**) Levels of transcripts coding for exosome subunits and DIS3 paralog DIS3L2. (**B**) Levels of mRNAs of proteins involved in RNA degradation, SKI complex subunits, and XRN1 and XRN2. (**C**) Levels of mRNAs coding for tRNA splicing factors and translation initiation factor eIF2α kinases. Blue bars, total mRNA; orange bars, mRNAs isolated from polysomes.

As pointed out above, the cytoplasmic RNases XRN1 and DIS3L2 had their expression increased upon infection. DIS3L2 has been characterized as an important 3’-5’ cytoplasmic exoribonuclease involved in mRNA decay [13, 14, 33, 34], and as shown here, has its expression increased at the translational level 6 hpi (Figs. 1B and 2A). The gene bank database indicates the expression of multiple DIS3L2 isoforms that can be detected by RT-PCR with common sets of primers. To identify the particular DIS3L2 isoforms expressed in macrophages, we performed RT-PCR with primers complementary to exons 8 and 17 (Suppl. Table 2) and amplified three DIS3L2 variants, allowing the identification of isoforms V2, X11 and X4 in three different murine cell lines, RAW 264.7, fibroblast L929 and myeloma-derived P3U1 (Fig. 3). Isoform V2 contains the RNA binding domains CSD1 and CSD2 at the N-terminus and S1 at the C-terminus, and the central exoribonuclease domain RNB, and we found it to be expressed in these three cell lines (Fig. 3B, upper band). This upper band may also correspond to variant V1 (Suppl. Fig. 1), but RT-PCR with primers complementary to exons 1 and 7 showed that only Δ6 is detected, which corresponds to V2 (data not shown). We also identified the variants X11 and X4 expressed in these cells (Fig. 3B, middle and lower bands, respectively). Additionally, the expression of these isoforms was analyzed in mouse peritoneal and embryonic stem cells (mESC), comparing with RAW 264.7. The results show that these variants are expressed in all three cells (Suppl. Fig. 1). The structure of mouse DIS3L2 has been determined, which shows its conservation relative to the yeast DIS3 ortholog Rrp44, despite the absence of the N-terminal PIN domain in DIS3L2 [17]. Interestingly, despite missing exons 13 or 14, respectively, isoforms X11 and X4 maintain the active site of the enzyme (of which residue D389 is part), although the RNB domain is shortened (Fig. 3).

**Figure 3.**
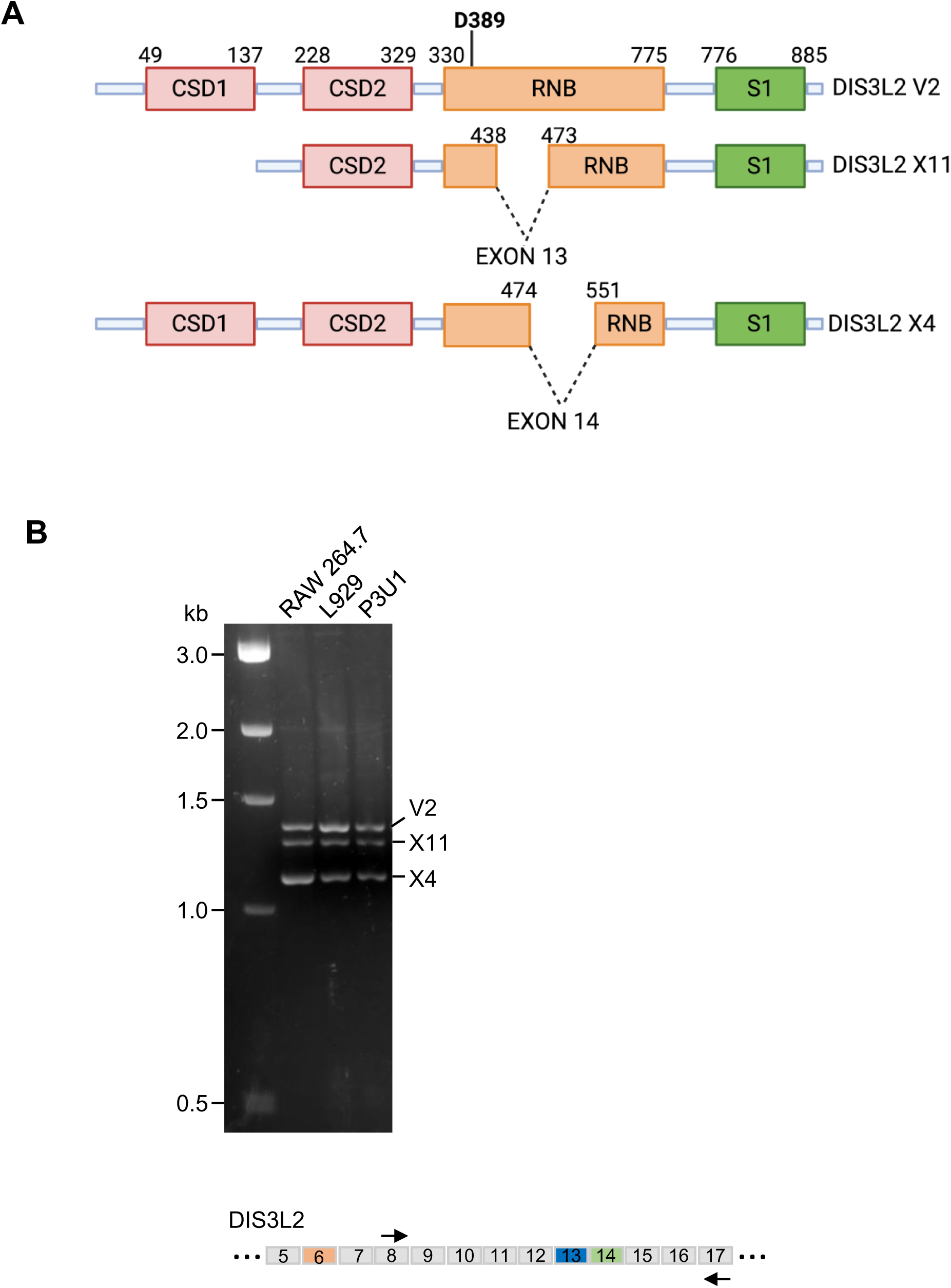
Comparison of the structure of the analyzed DIS3L2 isoforms. (**A**) Schematics of DIS3L2 isoforms. Numbers above indicate the amino acids residues in the full-length protein. D389 indicates the component of the DIS3L2 active site. (**B**) RT-PCR products, amplified with primers that anneal to DIS3L2 exons 8 and 17, show the expression of the expressed isoforms of DIS3L2, in mouse cell lines macrophage RAW 264.7, fibroblast L929, and myeloma-derived P3U1. Schematics of DIS3L2 cDNA, with the alternative exons 6, 13, and 14 highlighted in different colors is shown at the bottom.

The levels of expression of DIS3L2 isoforms were further assessed by RT-qPCR reactions with total RNA from three different mouse cell lines and specific pairs of primers (Suppl. Table 3). The results show little variation across cell lines of isoforms lacking, or containing exons 13, 14 and 6 (Fig. 4). However, the levels of isoforms display large differences in abundance in each cell line. Isoforms lacking exon13 (including X11) are present in very low levels, compared to isoforms containing exon13 (including V2 and X4). Isoforms lacking exon14 (including X4) are less abundant than those containing exon 14 (including V2 and X11). Isoforms lacking exon 6 (including V2, X4 and X11), on the other hand, show higher levels than those containing exon 6 (Fig. 4). Based on the comparison of the isoforms levels in these three cell lines, we can conclude that the abundance follows the order V2>X4>X11 (Fig. 4). These results suggest specific roles for these isoforms in the regulation of gene expression at posttranscriptional level.

**Figure 4.**
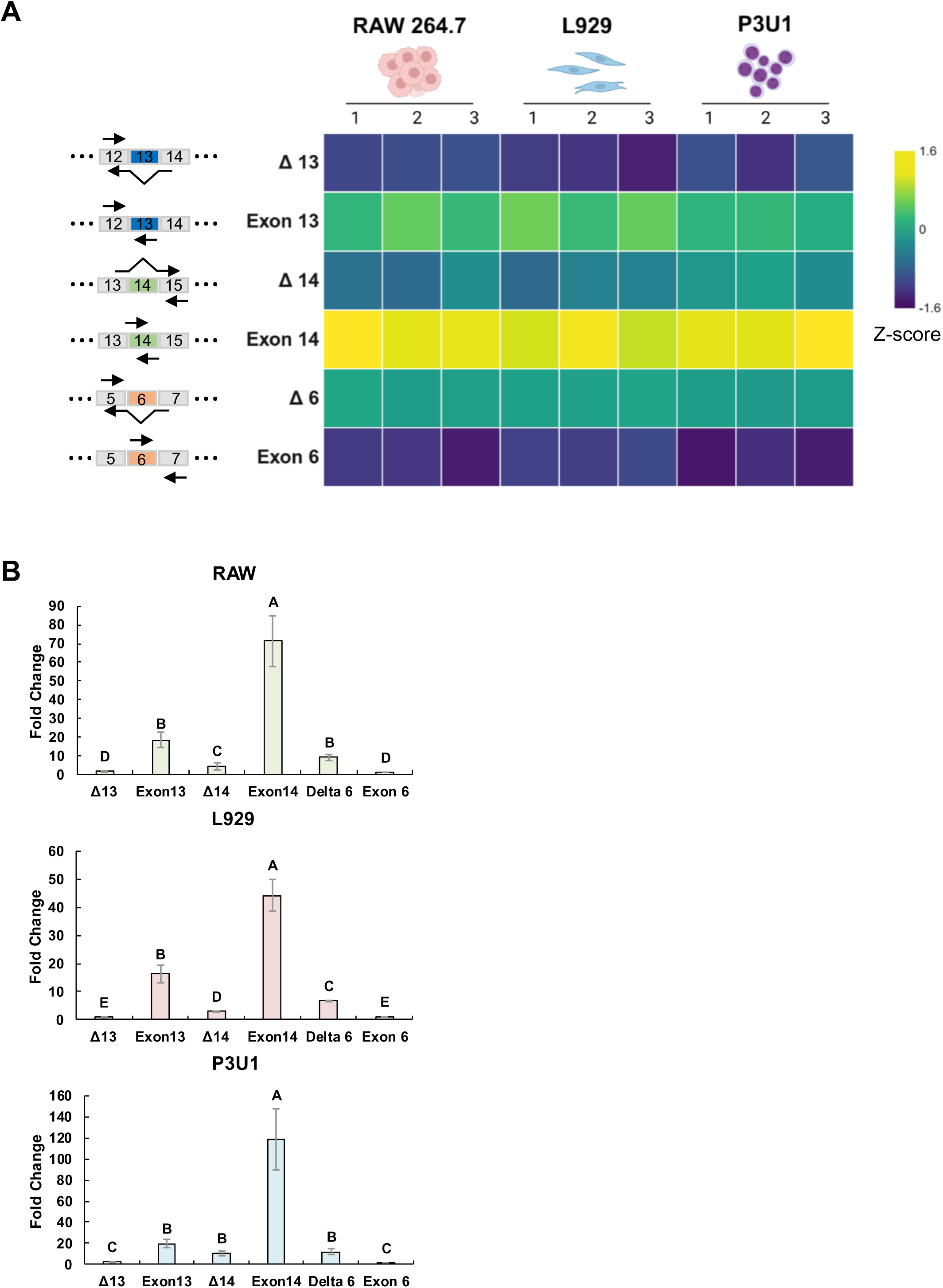
Relative expression of gene DIS3L2 isoforms in different mouse cell lines. (**A**) Heatmap of DIS3L2 isoforms expression in different cell lines determined by RT-qPCR. Primers were designed to detect the presence or absence of exon 13, 14, and 6, allowing the identification of the prevalence and abundance of the isoforms. Schematics of portions of DIS3L2 cDNA, with the alternative exons 6, 13, and 14 highlighted in different colors, with the respective primers used for the RT-PCR assays is shown on the left. (**B**) Comparison of expression levels of DIS3L2 in each cell line, as indicated. Ct values were normalized using the geometric mean of the reference genes ACT, GAPDH, and CYC1, and calibrated to the minimum value (exon 6). Fold change was calculated using the 2^−ΔΔCt^ method. Data are presented as the mean ± SEM from three biological replicates (each with three technical replicates). Statistical significance was determined by one-way ANOVA followed by Tukey’s post-hoc test. Different letters indicate significant differences between DIS3L2 isoforms (p < 0.05).

The observation that X11 and X4 have shortened RNB domains relative to V2, led to the question of whether these isoforms retain exonucleolytic activity. To test that, the three recombinant isoforms were purified from *E. coli* (Suppl. Fig. 2) and tested in RT-FeDEx assays [26, 27]. In this assay, a 5’ end FAM-labeled 30 nt single-stranded RNA (5’-FAM-RNA) and a single-stranded 17 nt DNA with a fluorophore bound its 3’ end (DNA-TAMRA-3’) are annealed and incubated with the proteins in an RT-PCR equipment for the detection of fluorescence throughout the reaction time. The FAM fluorescence is quenched by TAMRA, but as RNA degradation by the RNase proceeds, FAM is released, leading to an increase in the FAM fluorescence. The DNA oligo is complementary to the 5’-end portion of the RNA, such that the RNA contains a 3’ 13 nt single strand overhang that facilitates its 3’-5’ exonucleolytic degradation. These RNA degradation assays showed that all variants have exonucleolytic activity *in vitro*, but with differences in the efficiency of degradation (Fig. 5). As expected, higher fluorescence levels were emitted from samples incubated with V2 than with the other variants that have shortened RNB domains (Fig. 5A). X4 showed the lowest activity level, although still degrading RNA, despite missing part of the RNB domain. At the end of the RT-FeDEx reactions, samples were separated on denaturing polyacrylamide gels and fluorescence was detected on a Chemidoc MP equipment (BioRad) for analysis of the bands, which shows the degradation products of the reactions, comparing with RNaseA, and using the TAMRA-DNA, which is not degraded, as loading control (Fig. 5B). These results confirm that the three recombinant DIS3L2 isoforms are active for RNA degradation and their differences in activity might be due to alterations in their respective active sites. X11 has lost CSD1 domain, which, together with domains CSD2 and S1, control the access of the substrates to the active site funnel of DIS3L2 [17]. Therefore, it is possible that despite being shorter, the active site is more accessible to substrates in X11, allowing the RNA degradation by this variant. X4, on the other hand, contains CSD1 and CSD2, but has a different deletion in its RNB domain, which might compromise the conformation of the active site, affecting it negatively, as discussed below. X4 degrades RNA less efficiently, and as can be seen when merging the FAM and TAMRA signals, one of the degradation intermediates overlaps with the DNA band (indicated by an arrowhead in Fig 5B, upper panel), suggesting that X4 does not efficiently degrades double strand RNAs, as the other isoforms, resulting in lower concentrations of the final degradation product (Fig. 5B). Collectively, these results show that among the isoforms identified here, V2 has higher activity and expression levels than X11 and X4, X11 is the least abundant, but is almost as efficient as V2 in RNA degradation, while X4 has intermediate abundance and is the least efficient in degrading dsRNAs (Figs. 4 and 5).

**Figure 5.**
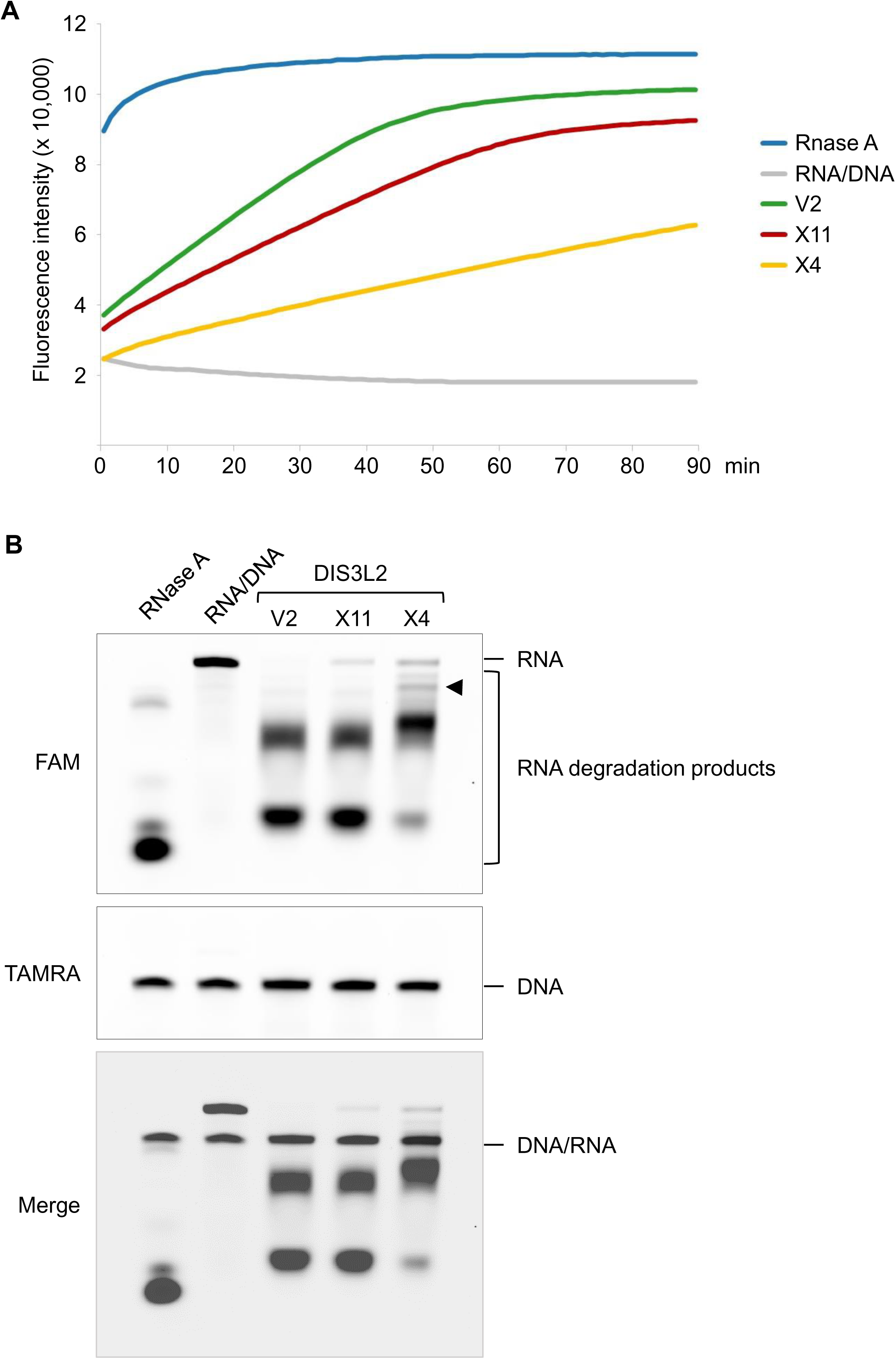
RT-FeDEx assay to analyze exonucleolytic activity of DIS3L2 isoforms. 5′-FAM-labeled RNA substrate annealed to a complementary single stranded 3′-TAMRA-labeled DNA was incubated at 25 °C in the presence of the purified recombinant DIS3L2 isoforms in a RT-PCR. The close proximity of TAMRA quenches FAM fluorescence, but once the RNA subtract is hydrolyzed, FAM fluorescence is detected in real time. (**A**) Plot of the fluorescence detected during the reaction. Values represent the absolute fluorescence emitted by FAM at the indicated time point. Data are from three technical replicates and bars represent standard deviation. Samples containing only RNA or the hybrid molecule RNA/DNA with no protein added, and incubation with RNaseA were used as controls. (**B**) After completion of the RT-FeDEx reaction, samples were separated on denaturing polyacrylamide gels and visualized on a Chemidoc MP (BioRad). Full length RNA used as substrate and the hydrolysis products of DIS3L2 isoforms and RNaseA are indicated on the right. Full length DNA is shown on the lower panel. Arrowhead indicates RNA band overlapping with DNA.

Due to their differences in the RNA binding domains content, we also tested the ability of the DIS3L2 variants to bind RNA *in vitro*. The same 5’-FAM labeled RNA used in the RT-FeDEx experiments was incubated with different concentrations of the recombinant proteins for electrophoresis mobility shift assays (EMSA). After incubation at 30°C for 30 min in the presence of EDTA, samples were separated by native polyacrylamide gel electrophoresis (PAGE) and visualized on a Chemidoc MP equipment (BioRad). The results show that the three isoforms bind RNA with similar affinity (Fig. 6).These results are very interesting, especially because X11 is missing the CSD1 RNA binding domain, although it still contains CSD2 and S1 domains, which might be sufficient for binding RNA in a sequence-independent manner.

**Figure 6.**
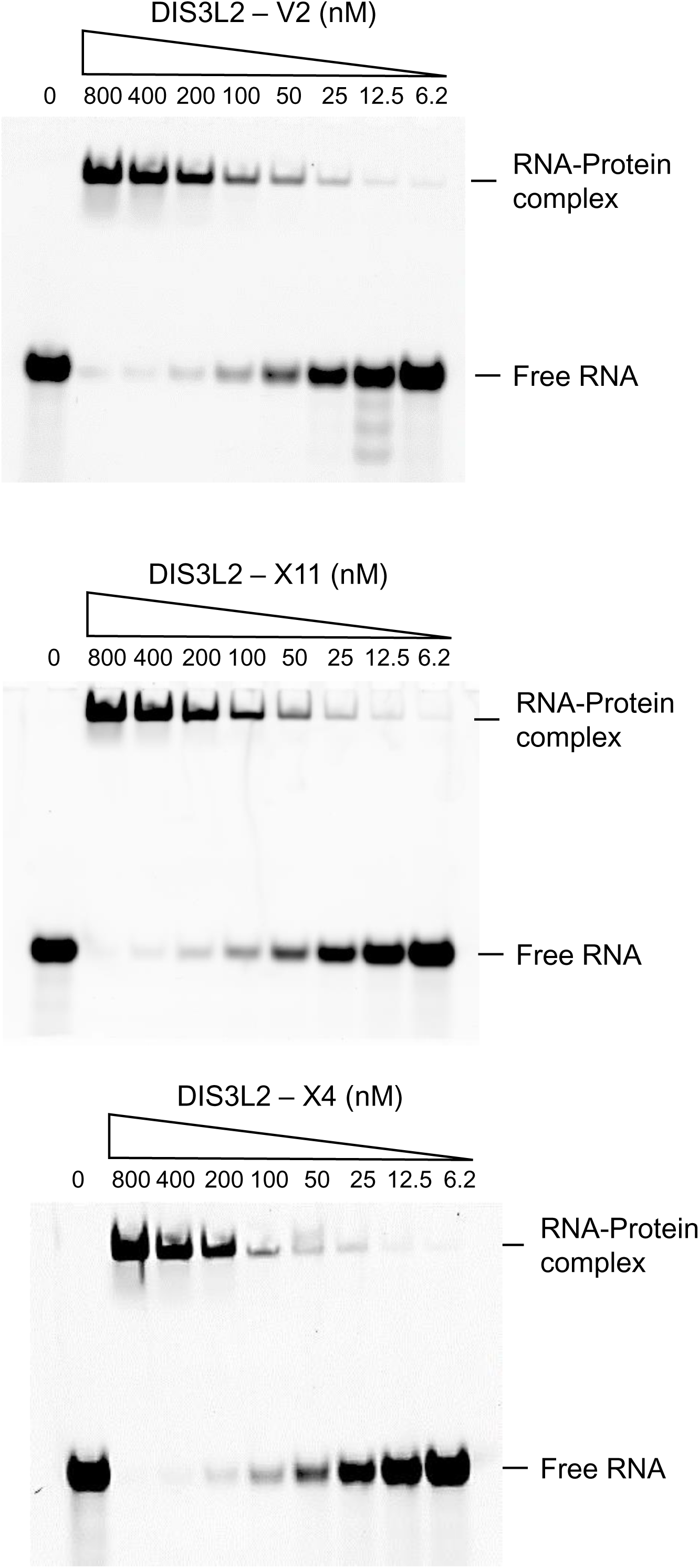
RNA-binding ability of Dis3L2 isoforms. EMSA was performed by incubating 1 pmol of 5’ pmol of-FAM-RNA substrate for 30 min at 30°C with decreasing concentrations of each Dis3L2 variant (indicated on the top): 0 nM (Control), 800 through 6.2 nM. Reaction products were resolved in a 12% native gel.

To analyze these isoforms in more detail, we next used AlphaFold3 to predict the structure of these proteins, comparing it with the crystallographic structure of mDIS3L2 [17]. The structure modeling shows that despite their deletions in the RNB domain, X11 and X4 maintain the general structure of DIS3L2, including the catalytic site coordinating a Mg^2+^ ion (Fig. 7), in agreement with the RNA degradation assay described above. The deletion of the region comprising amino acid residues 475-550 results in the absence of two alpha helices and a strand of a beta sheet in DIS3L2-X4, close to the active site, which does not compromise RNA binding of this isoform, but can affect its RNase activity (Fig. 7A), nicely corroborating the activity assays described above. DIS3L2-X11 has a deletion in the region containing amino acid residues 439-472, resulting in the absence of a beta strand and a short alpha helix portion of the RNB domain, which does not affect the active site (Fig. 7B). These comparisons can be better visualized by careful analysis of the alpha helix that is missing in X4, but is present in X11, where Tyr492 is located. Its side chain can directly interact with the RNA base upstream of the 3’-end and may be involved in directing the RNA to the active site (Fig. 8). Combined with the activity assays, these structural analyses confirm that all identified isoforms are active RNases, but show differences that may be important for playing particular roles in posttranscriptional control of gene expression.

**Figure 7.**
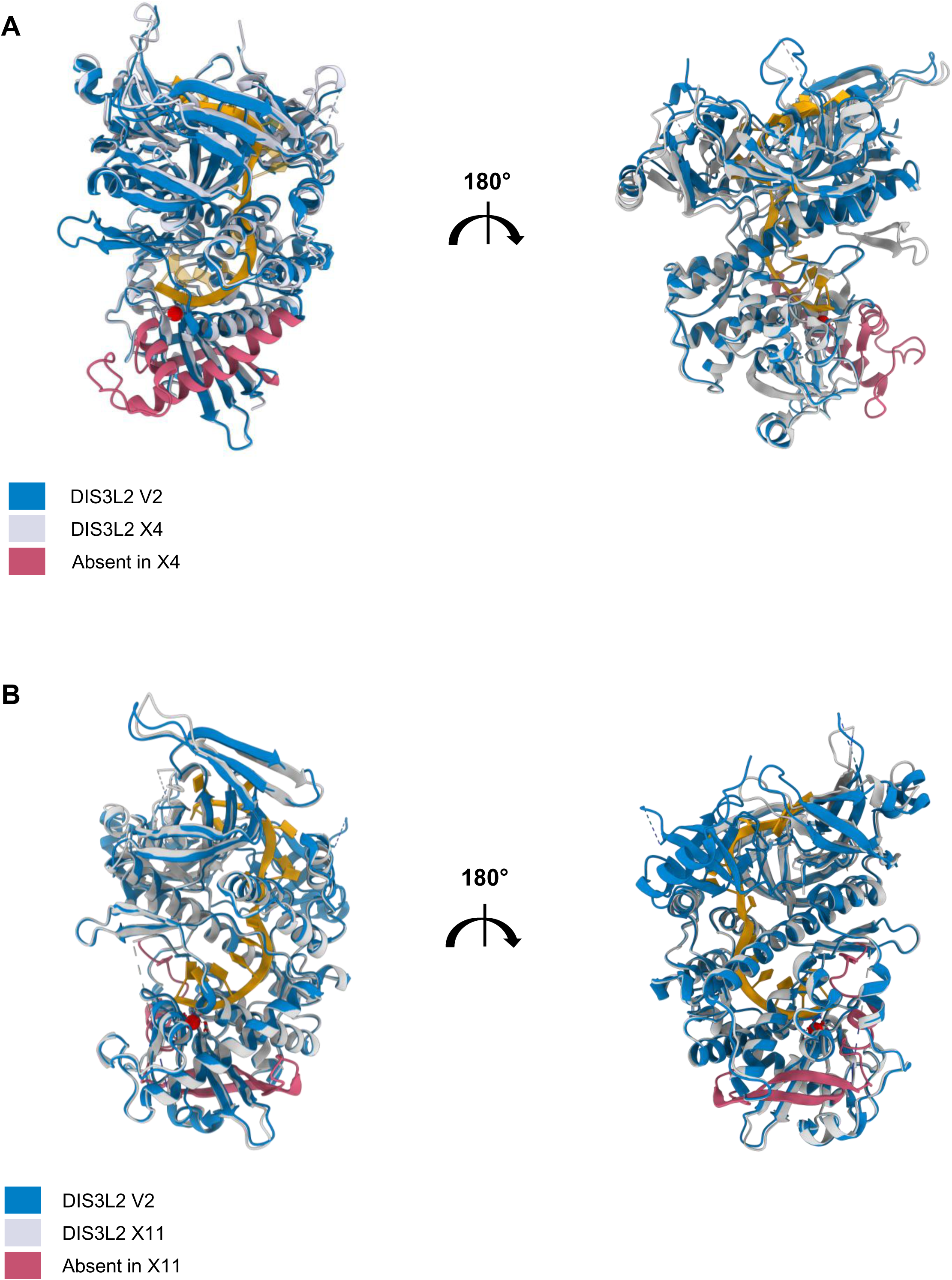
Structure modeling of Dis3L2 isoforms. Alphafold prediction of the structures of DIS3L2 isoforms. (**A**) Comparison of V2 (blue) with X4 (gray). An alpha helix close to active site (pink) is missing in X4. RNA substrate is represented in orange. Mg^2+^ is shown in red. (**B**) Comparison of V2 (blue) with X11 (gray). A beta sheet and linker (pink) is missing in X11. Based on crystal structure of mDIS3L2 (PDB: 4PMW).

**Figure 8.**
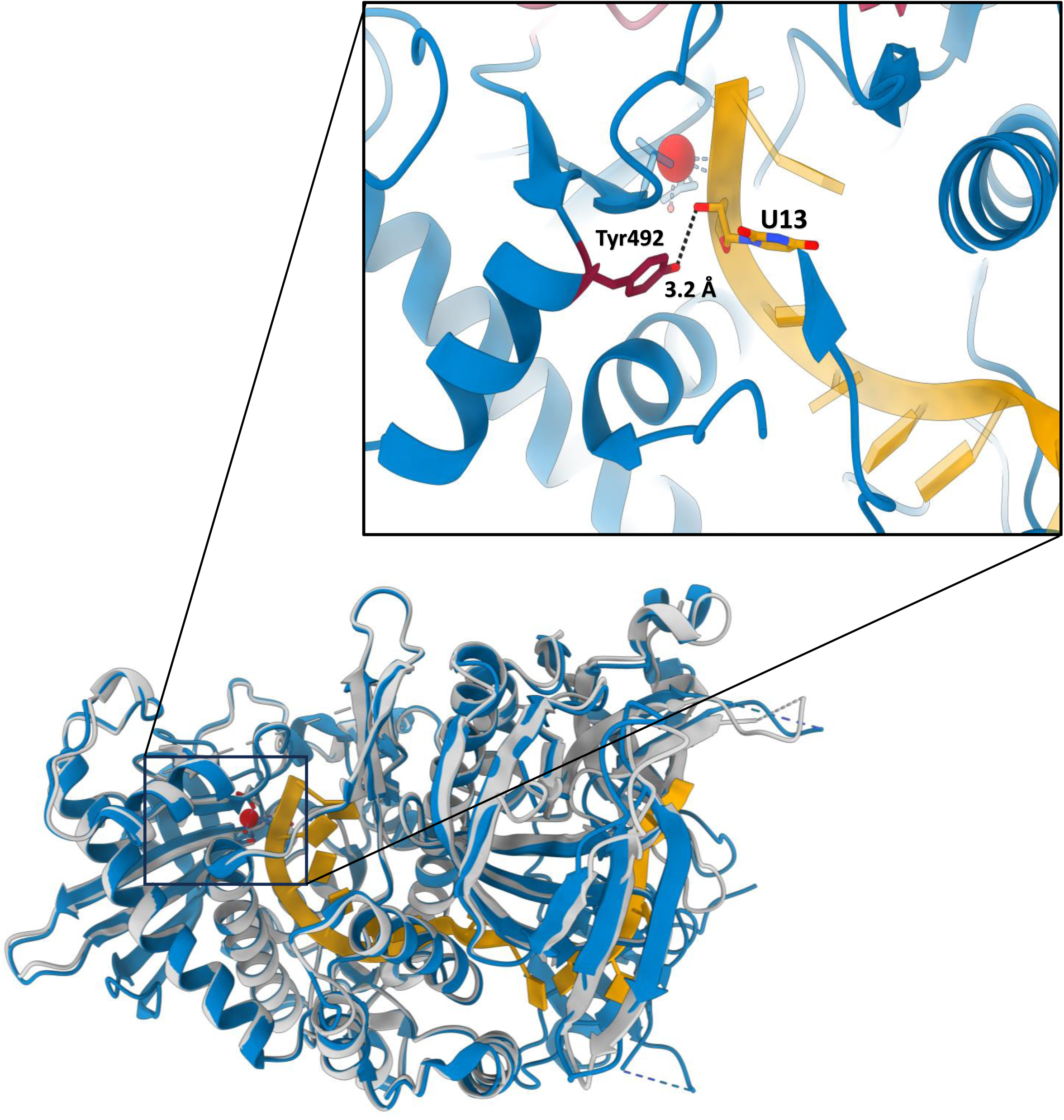
Structure modeling of Dis3L2-X11 isoform interaction with RNA. Alphafold prediction of the structure of DIS3L2 V2 and X11 isoforms bound to RNA to detail Tyr492 of an alpha helix present in both isoforms directly binding RNA (orange). Mg^2+^ is shown in red. Based on crystal structure of mDIS3L2 (PDB: 4PMW).

To better understand the role of DIS3L2 in gene expression regulation, we performed pulldown experiments to identify proteins that might recruit or control the activity of DIS3L2 on cytoplasmic RNA degradation. These experiments were carried out by incubating purified recombinant DIS3L2 isoforms and macrophage extracts. The isolated proteins were identified by mass spectrometry, which showed that 427 were pulled down with the three isoforms, while different number of proteins interacted exclusively with each isoform (Suppl. Fig. 3; Suppl. Table 4). Gene ontology analysis shows that the enriched categories of proteins interacting with DIS3L2 V2 are related to mRNA processing, and translation control (Suppl. Fig. 4A). Interestingly, the most enriched category of proteins co-purified with V2 is L13a-mediated translational silencing of ceruloplasmin expression. L13a is a ribosomal protein that has been shown to be released from the 60S subunit upon phosphorylation by proinflammatory interferon γ-induced cytokine in macrophage, and to join the inhibitor of translation complex GAIT [35]. The hypothesis of DIS3L2 interacting with L13a is reinforced by the identification of the association of DIS3L2 with polysomes [13], therefore allowing the control of gene expression by translation repression and mRNA degradation. In addition, among the most enriched proteins pulled-down with V2 is the ribosomal protein RPL24, RNA helicases, translation control factors, and protein components of P-bodies (Fig. 9A; Suppl. Table 5), further confirming DISL2-V2 interaction with ribosomes and involvement with RNA degradation. DIS3L2 has also been shown to be involved in quality control of snRNAs [36], which further validates our results, since we also isolated proteins of the mRNA processing and splicing regulation pathways (Fig. 9A; Suppl. Fig. 4A).

**Figure 9.**
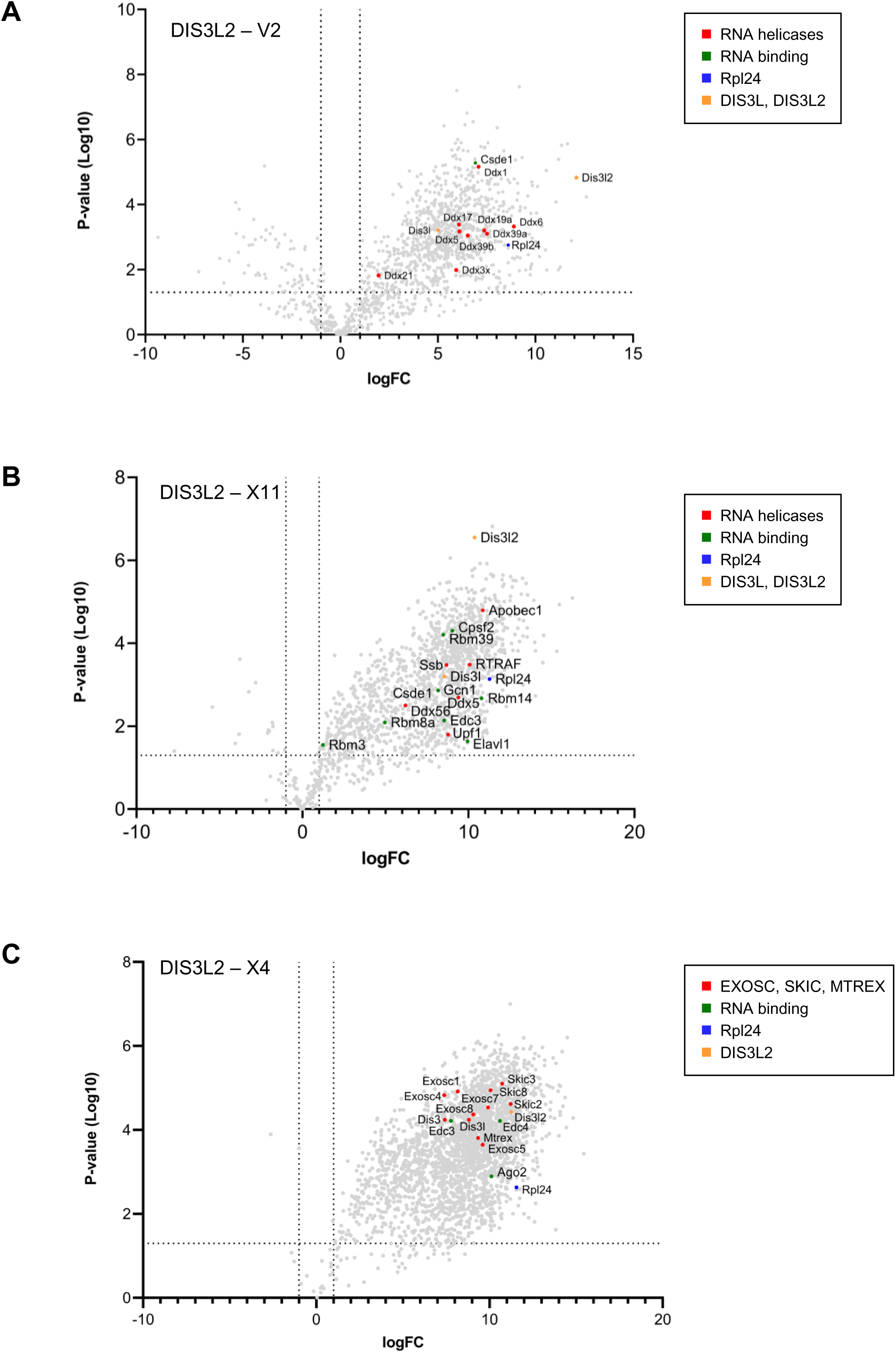
Proteomics analysis of macrophage proteins pulled-down with DIS3L2 isoforms. Volcano plot comparing the two conditions (pull-down with resin-bound DIS3L2 vs resin) indicating the different proteins interacting with the isoforms. (**A**) DIS3L2-V2; (**B**) DIS3L2-X11; (**C**) DIS3L2-X4. Proteins with related functions are highlighted in similar colors.

DIS3L2 isoforms X11 and X4 were also used in pulldown experiments and, interestingly, showed different interactome profiles, although both still interact with Rpl24 and other proteins involved in translation and mRNA metabolism. The most striking difference is that X11 and X4 do not pulldown proteins from the L13a-mediated translational control pathway (Suppl. Fig. 4A), suggesting specific mechanisms of control of gene expression. The most enriched category of proteins co-purifying with X11 is related to ribosomal subunits joining, suggesting a role in translation control, further confirming the ribosome interaction of DIS3L2, and specific functions for each isoform. In addition to Rpl24, proteins interacting with X11 are known for being involved in RNA metabolism, such as the cytidine deaminase APOBEC1, cleavage and polyadenylation factor CPSF2, cap-binding complex subunit RTRAF, and non-sense mediated decay (NMD) factor UPF1 (Fig. 9B).

DIS3L2-X4 also pulled-down Rpl24 and RNA metabolism-related proteins, underscoring the specificity of the results obtained here (Suppl. Fig. 4A). Interestingly, X4 pulled-down many exosome subunits, subunits of the SKI complex (a cytoplasmic exosome cofactor), the RNA helicase MTREX, and Argonaute 2 protein. The interactions identified here are not necessarily direct, but the exosome subunits being pulled-down with X4 is striking because DIS3L2 is known for not interacting with the exosome since it lacks the N-terminal portion of DIS3 that is responsible for this interaction. DIS3L2-X4 interactome suggests the involvement of this isoform in RNA degradation processes in which the exosome and the SKI complex also participate. The results shown here underscore the importance of DIS3L2 in the control of cytoplasmic RNA metabolism and highlights the specific roles its isoforms might play in this pathway.

## Discussion

RNases are an important factor for posttranscriptional regulation of gene expression in eukaryotes. In the cytoplasm, the 3’-5’ exoribonuclease RNA exosome, for instance, is involved in mRNA decay pathways, many of them dependent on translation [37]. Here, we set out to analyze the role the 3’-5’ RNases RNA exosome and DIS3L2 might play in the control of gene expression in murine macrophage and investigate whether it is modulated by infection with *Leishmania major*.

As shown here, the host cell responds to infection by up- or downregulating different groups of genes. The major changes occur on genes of the protein transport pathway, which are highly upregulated 6 hpi, and genes of the ribosomal biogenesis pathway, which includes the exosome. Interestingly, the core exosome subunits are downregulated upon infection, whereas the catalytically active subunits, DIS3, DIS3L and EXOSC10, are upregulated. These intriguing results suggest that the core exosome might associate dynamically with the active subunits and recycle at the end of each reaction. This could be explained by the evidence supporting the role of the core in the recruitment and regulation of exosome activity. Corroborating this hypothesis, cytoplasmic exosome cofactor SKI complex is also upregulated upon infection, as well as the exonucleases XRN1, XRN2, and DIS3L2. These results indicate that the host cell responds rapidly to infection at the posttranscriptional level in the first hours, and the response decreases over time, confirming previous data [21, 38].

DIS3L2 is a cytoplasmic DIS3 paralog that has 3’-5’ exonuclease activity but does not associate with the exosome. After detecting DIS3L2 upregulation upon infection, we identified the DIS3L2 isoforms expressed in macrophage RAW 264.7, lacking exon 6 (V2) and either exon 13 (variant X11), or exon 14 (variant X4). Here we show that these three isoforms are expressed in macrophage and other cell lines. Importantly, we show that these three variants have exoribonuclease activity and bind RNA *in vitro*. DIS3L2 binds RNA through S1 and CSD domains and is directly involved in cytoplasmic decay of various classes of RNAs, such as mRNAs, microRNAs and other noncoding RNAs [13, 17, 39–41]. By comparing the structures of the DIS3L2 isoforms identified here, we show that despite missing CSD1, DIS3L2-X11 efficiently binds RNA and shows exonucleolytic activity very similar to isoform DIS3L2-V2. DIS3L2-X4, on the other hand, degrades RNA less efficiently probably because it lacks an alpha helix that may participate in directing the substrate to the catalytic site.

Contrary to the cytoplasmic exosome subunit DIS3L, DIS3L2 can degrade structured RNAs, highlighting its importance for the control of gene expression [17, 34]. In addition, DIS3L2 has been shown to affect alternative splicing [42]. It is therefore important to determine how DIS3L2 is recruited to the target RNAs for degradation. With this aim, we performed protein pull-down experiments with recombinant DIS3L2 variants and macrophage cell extracts and identified proteins that might interact with DIS3L2, which may recruit or control its activity. Interestingly, all three DIS3L2 isoforms co-purified ribosomal protein Rpl24, suggesting a role for DIS3L2 in the control of ribosome associated mRNAs. The identification of DIS3L2 bound to ribosomes [13] underscores the validity of the results shown here. Importantly, although all proteins identified in the pull-down experiments have functions related to RNA metabolism, each isoform showed a particular interactome. DIS3L2-V2 co-purified many RNA helicases, translation control factors and P-bodies proteins. DIS3L2 isoforms X11 and X4 also pulled down ribosome-associated proteins, most of them involved in translational control or protein folding, strengthening the hypothesis of ribosome association for efficient expression control. The two exonucleolytic RNases with complementary activities DIS3L2 (3’-5’) and XRN1 (5’-3’) have been shown to associate with each other in an RNA-dependent manner, to affect cell proliferation and apoptosis, and mutations in their genes are associated with human diseases such as Perlman syndrome and some types of cancer [13, 43]. Accordingly, we show that DIS3L2-X11 pulled down NMD factor UPF1, and further confirming our results, DIS3L2 and XRN1 have been shown to participate in nonsense-mediated decay (NMD), a process of mRNA degradation dependent on translation [44]. Surprisingly, DIS3L2-X4 pulled down exosome subunits, as well as exosome cofactors, such as subunits of the SKI complex, and other RNases, such as EDC3/4 and Ago2. These results strongly suggest the participation of DIS3L2 in RNA degradation pathways in which the exosome and other RNases are also involved. The results shown here strongly indicate a particular function for each DIS3L2 isoform and the proteins identified here as DIS3L2 isoforms interactors provide important information towards the characterization of the mechanisms regulating their functions. Further studies will show whether the human isoforms of DIS3L2 also have specialized functions.

## Authors contributions

LPPC, CCO, HG planned experiments; LPPC, BRSQ, KT, VGN, LHSM, BAU performed experiments; LPPC, BRSQ, KT, VGN, HG, CCO analyzed data; HG, CCO acquired funding; CCO wrote the paper; all authors reviewed, edited and approved the text.

## Acknowledgements

We are grateful to all members of the Oliveira and Goto laboratories, especially to Eduardo M. Ramos-Sanchez and Fernando A. Gonzales-Zubiate, for ideas, help, reagents, and discussion. We are especially thankful to Mauricio da Silva Baptista for the use of his laboratorýs cell culture infrastructure and reagents. This research used facilities of the Brazilian Biosciences National Laboratory (LNBio-CNPEM). The NuTeP staff is acknowledged for the assistance during the experiments (MAS-20252621).

## Funding

This work was supported by grants from Fundação de Amparo à Pesquisa do Estado de São Paulo (FAPESP -20/00901-1, 22/08812-3 to CCO), and in part by the Coordenação de Aperfeiçoamento de Pessoal de Nível Superior (CAPES). LPPC was supported by a CNPq (Conselho Nacional de Desenvolvimento Científico e Tecnológico) fellowship. BRSQ, VGN, KT, and LHSM were supported by FAPESP fellowships (23/06344-5, 22/00071-4, 24/02036-7, 18/17562-5, respectively).

## Notes

Conflicts of Interest: The authors declare no conflicts of interest.

https://www.ebi.ac.uk/pride/

https://www.ncbi.nlm.nih.gov/sra

### Competing Interest Statement

The authors have declared no competing interest.

