## Supplementary material for "Identification and functional characterization of novel murine DIS3L2 isoforms": Supplental Figures

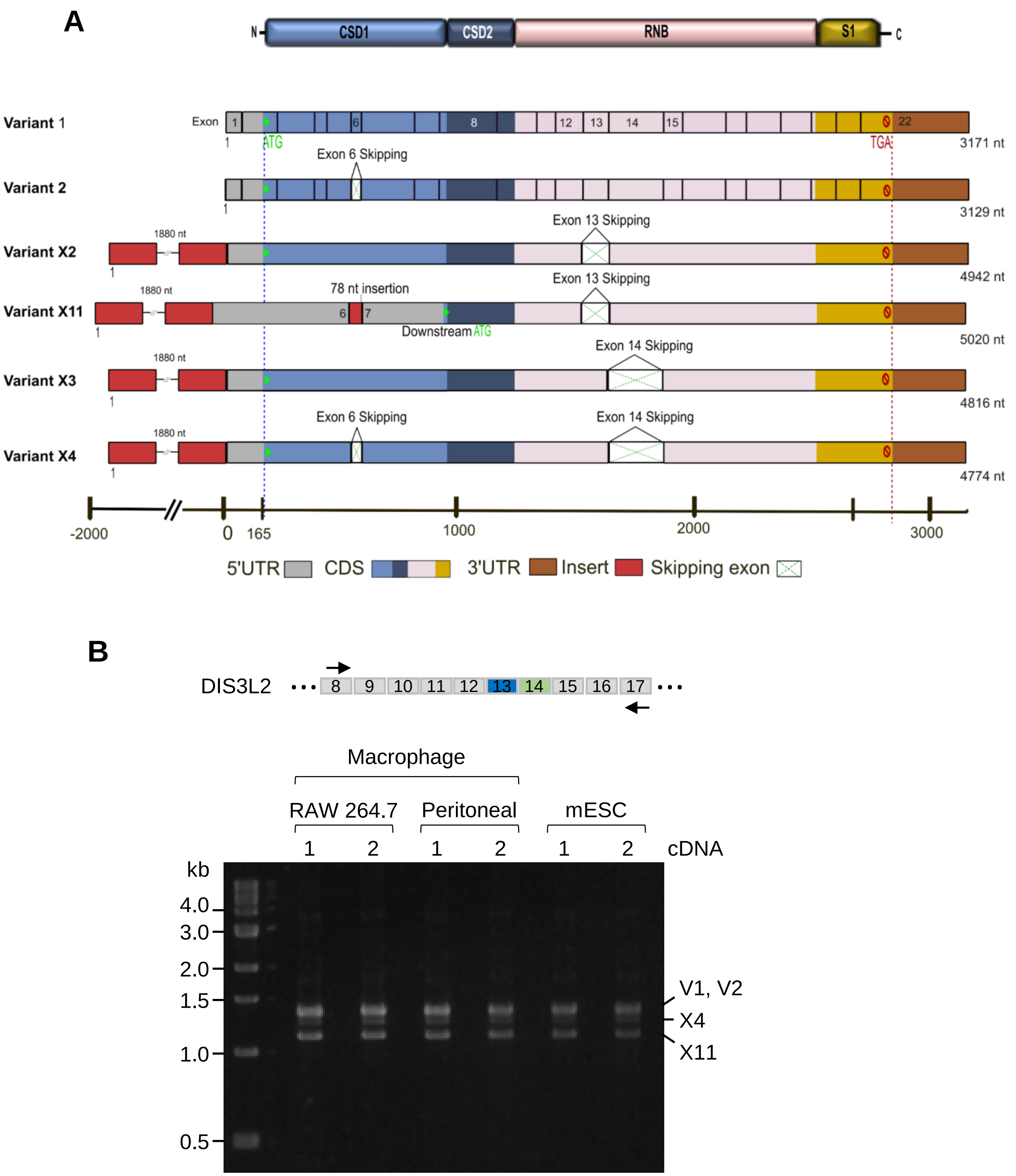


**Suppl. Figure 1.** ***DIS3L2 isoforms are expressed in mouse macrophages and ESC.*** (**A**) Schematics of DIS3L2 isoforms. Numbers above indicate the amino acids residues in the full length protein. D389 indicates the component of the DIS3L2 active site. (**B**) RT-PCR products, amplified with primers that anneal to DIS3L2 exons 8 and 17, show the expression of the indicated DIS3L2 isoforms in macrophage cell line Raw 264.7, mouse peritoneal macrophage cells and embryonic stem cells.

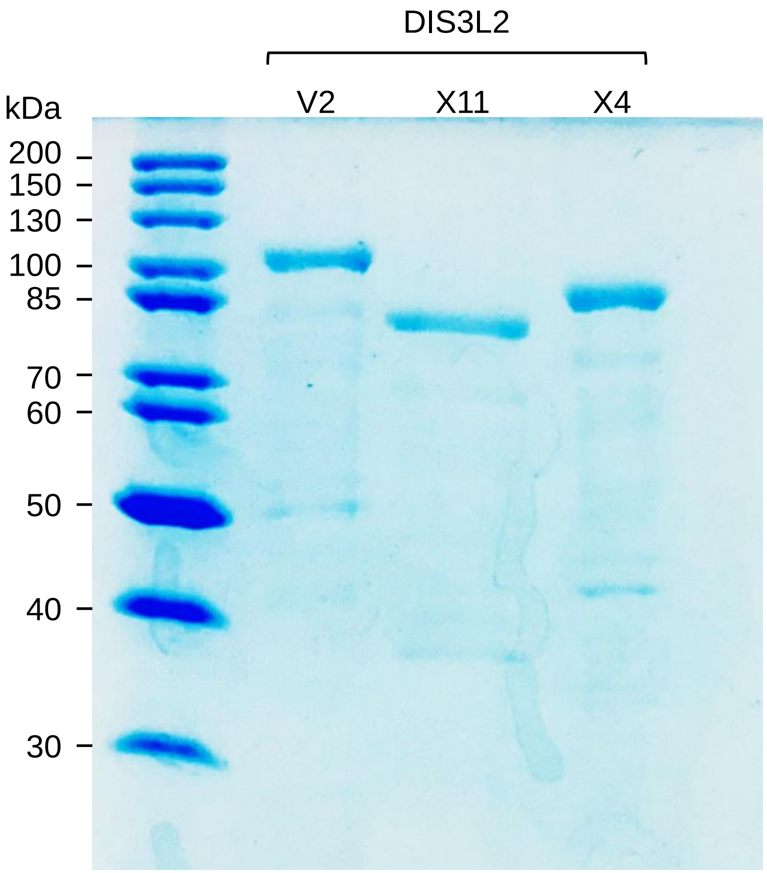


**Suppl. Figure 2. Purification of recombinant DIS3L2 isoforms.** SDS-PAGE of purified DIS3L2 isoforms stained with coommassie blue show the purified recombinant proteins used in RT-FedEx activity assays.

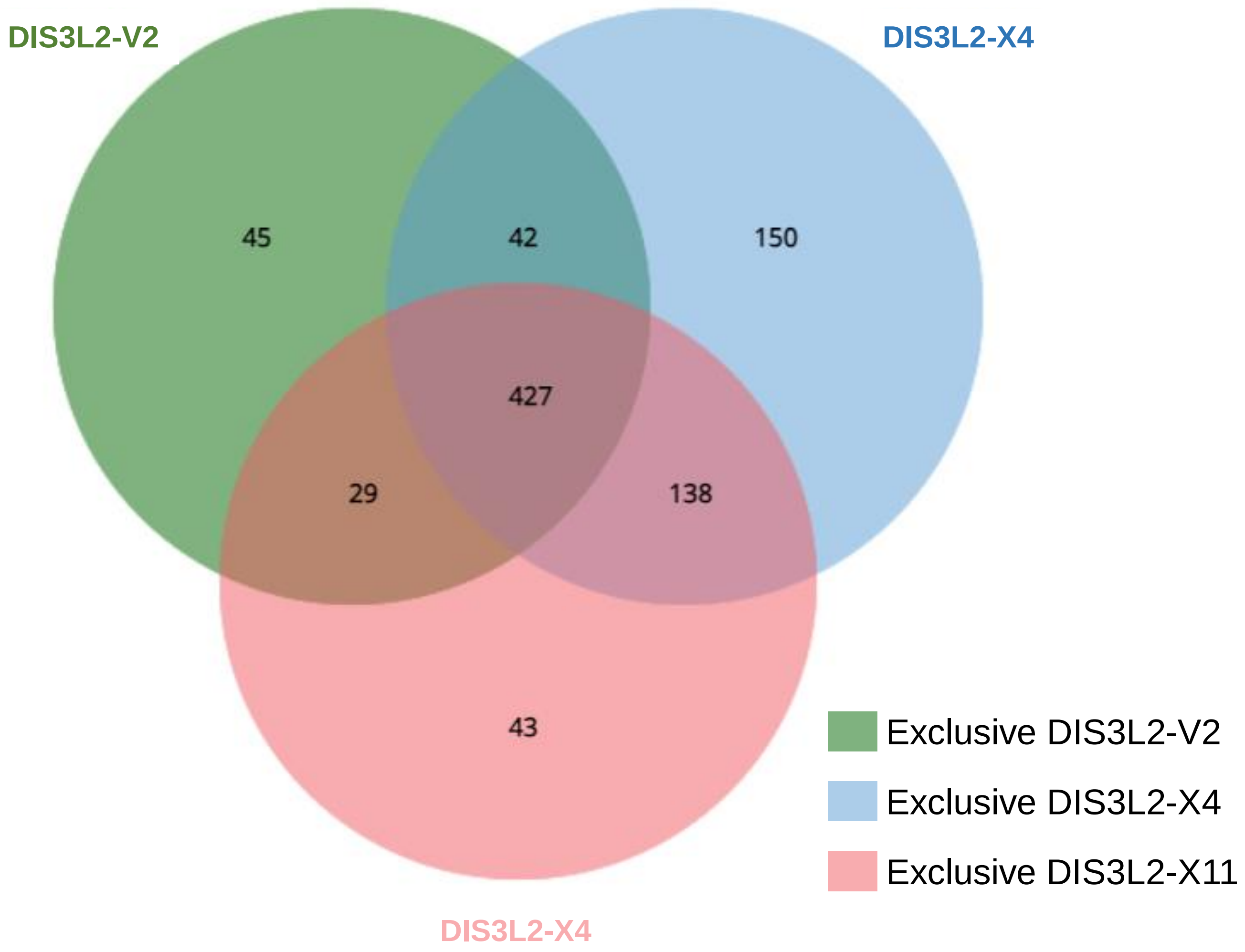


**Suppl. Figure 3. Pull down of macrophages proteins with purified recombinant isoforms DIS3L2-V2, DIS3L2-X11 and DIS3L2-X4.** Venn diagram of the proteomics analysis of proteins pulled-down with DIS3L2 isoforms. Numbers indicate the proteins that were co-purified exclusively with each of he isoforms, or in common with more than one isoform, indicated by different colors.

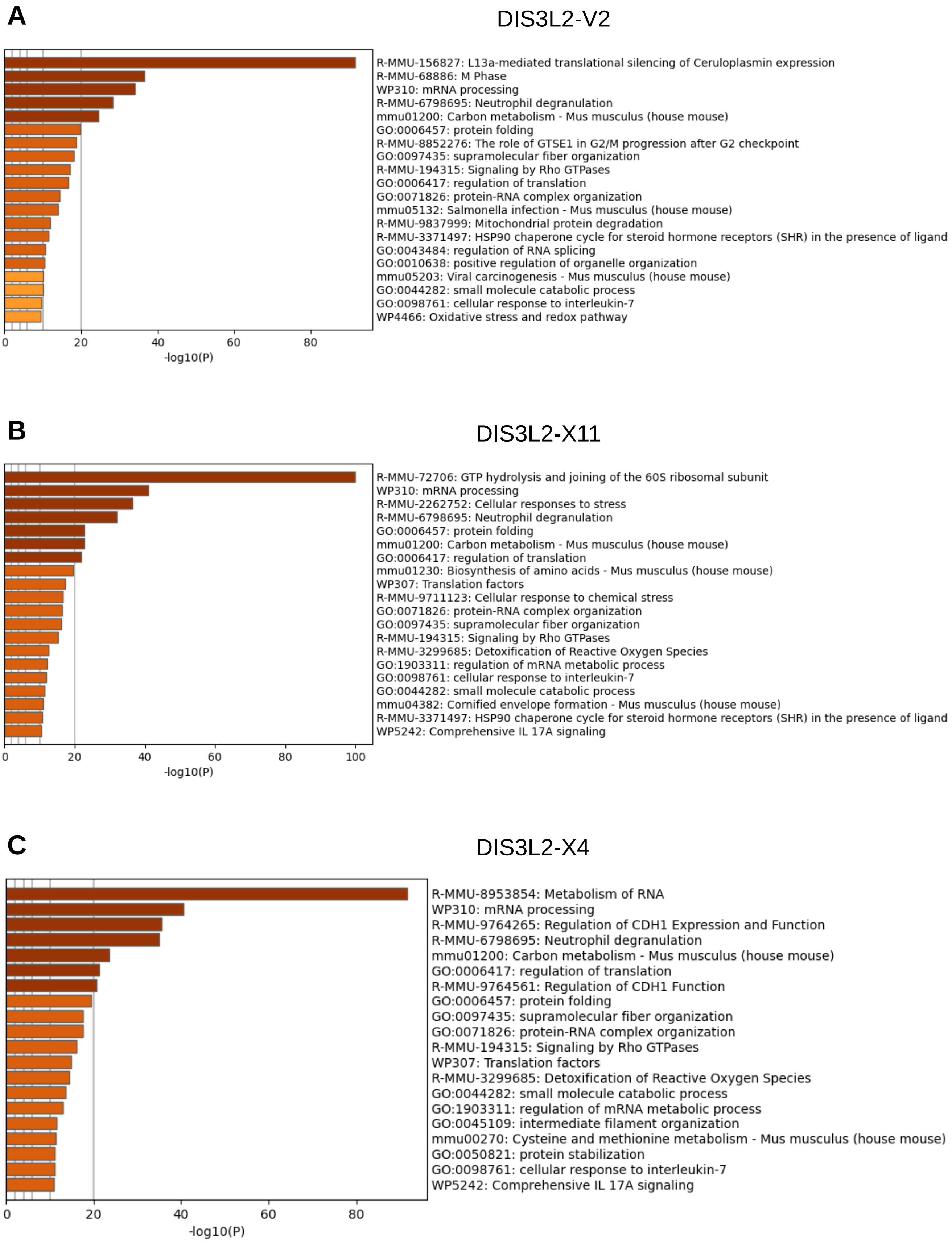


**Suppl. Figure 4. Pull down of macrophages proteins with purified recombinant isoforms DIS3L2-V2, DIS3L2-X11 and DIS3L2-X4.** Gene onthology enrichment according to biological processes. Proteins co-purified with DIS3L2-V2 (**A**), DIS3L2-X11 (**B**), and DIS3L2-X4(**C**).
